# A Single-Cell Approach Reveals Variation in Cellular Phage-Producing Capacities

**DOI:** 10.1101/2021.10.19.465070

**Authors:** Sherin Kannoly, Gabriella Oken, Jon Shadan, David Musheyev, Kevin Singh, Abhyudai Singh, John J. Dennehy

## Abstract

Virus burst size, a component of viral fitness, is the average number of viral particles released from a single infected cell. In this study, we estimated bacteriophage lambda (λ) burst size mean and distribution at different lysis times. To estimate phage λ burst sizes at single-cell level, we employed a lysis-deficient *E. coli* lysogen, which allowed chemical lysis at desired times after the induction of lytic cycle. Induced cultures of *E. coli* lysogen were diluted and aliquoted into the wells of 96-well plates. A high dilution rate results in mostly empty wells and minimizes the probability of having multiple cells in wells that do receive cells. Burst size was estimated by titering single-cell lysates obtained after chemical lysis at desired times. Our data shows that the viral burst size initially increases exponentially with the lysis time, and then saturates at longer lysis times. We also demonstrate that cell-to-cell variation or “noise” in lysis timing does not significantly contribute to the burst size noise. The burst size noise remains constant with increasing mean burst size. The most likely explanation for the experimentally observed constant burst size noise is that cell-to-cell differences in burst size originate from differences in cellular capacity to produce phages. The mean burst size measured at different lysis times is positively correlated to cell volume, which may determine the cellular phage production capacity. However, experiments controlling for cell size indicates that there are other factors in addition to cell size that determine this cellular capacity.

**ARTICLE IMPORTANCE:** Phages produce offspring by hijacking a cell’s replicative machinery. Previously, it was noted that the variation in the number of phages produced by single infected cells far exceeded cell size variation. It has been hypothesized that this variation is a consequence of variation in the timing of host cell lysis. Here we show that cell-to-cell variation in lysis timing does not significantly contribute to the burst size variation. We suggest that the constant burst size variation across different host lysis times results from cell-to-cell differences in capacity to produce phages. We find that the mean burst size measured at different lysis times is positively correlated to cell volume, which may determine the cellular phage production capacity. However, experiments controlling for cell size indicates that there are other factors in addition to cell size that determine this cellular capacity.

## INTRODUCTION

As part of their life cycle, bacteriophages (phages) lyse host cells after assembling progeny virions. Burst size, which is defined as number of virions produced per infected host cell, and lysis time, which is defined as time elapsed between infection and lysis, are key traits that affect phage fitness. Generally, burst size is thought to be positively correlated to the lysis time–a longer lysis time will result in a higher burst size (Hutchison and Sinsheimer, 1966; Josslin, 1970; Reader and Siminovitch, 1971; Wang, Dykhuizen and Slobodkin, 1996). As phenotypic traits, burst size and lysis time are akin to fecundity and generation time of an organism and affects host-virus population dynamics.

Ellis and Delbrück developed the one-step growth curve to estimate the burst size and lysis time of a coliphage in liquid culture (Ellis and Delbrück, 1939). Since then, the one-step growth curve has been extensively used in phage biology to characterize phage life history traits (Doermann, 1952; Reilly and Spizizen, 1965; Fischetti, Barron and Zabriskie, 1967; Pollard and Tilberg, 1972; Rosario and Drake, 1990; Herrero *et al*., 1994; Wang, 2006; García *et al*., 2008; Nabergoj, Modic and Podgornik, 2018). Burnet invented the first method for estimating the single-cell burst size using phages, which was further modified by others to obtain distributions of phage burst sizes (Burnet, 1929; Delbrück, 1945).

Phage lambda (λ) has been extensively used as a model system to study lysis time, burst size, and their effects on phage fitness (Wang, 2006; Dennehy and Wang, 2011; Singh and Dennehy, 2014; Ghusinga, Dennehy and Singh, 2017; Kannoly *et al*., 2020). Following the induction of the lytic program, λ lysis genes *S*, *R*, *Rz*, and *Rz1* are expressed from the late promoter, *pR*’. The *S* gene has a dual-start motif and encodes both holin and its inhibitor, antiholin (Cahill and Young, 2019). Holin dimers accumulate in the inner membrane and reach a critical threshold concentration whereupon they form a micron-sized hole. This membrane lesion allows product of the *R* gene, endolysin, to access the periplasmic space and degrade the cell wall. Subsequently, the spanins, *Rz* and *Rz1*, disrupt the outer membrane and phage progeny are released into the surrounding medium (Cahill and Young, 2019).

In a previous study, we used a panel of phage λ holin mutants that varied in their lysis times to show that the cell-to-cell variation or “noise” in lysis timing takes a concave-up shape with respect to lysis time, suggesting an optimal lysis time that minimizes noise (Kannoly *et al*., 2020). A theoretical approach further suggests that the noise in lysis timing is minimized when both holin and its antagonist antiholin are expressed at an optimal ratio (Dey *et al*., 2020). In a follow up study, we elucidated the biological significance of minimizing the noise in lysis timing and demonstrated that there exists a range of optimal lysis times where phage fitness is maximized in a quasi-continuous culture (Kannoly, Singh and Dennehy, 2020). Encouraged by these results, we further explored the effect of lysis time on burst size. The conventional method only estimates the burst size values averaged across a phage population; therefore, it is not suitable to measure cell-to-cell variation (noise) in burst size. In this study, we elucidated the effects of lysis time on burst size using single-cell assays. To estimate phage λ burst sizes at single-cell level, we employed a lysis-deficient *E. coli* lysogen, which allowed lysis at desired times after the induction of lytic cycle. We modified the method used by Delbrück (Delbrück, 1945) to estimate the individual burst sizes of a large number of cells at different lysis times.

## MATERIALS AND METHODS

### Bacterial Strains

| Strain | Genotype | Source |
| --- | --- | --- |
| CGSC#: 6152<sup>a</sup> | *E. coli* MC4100 (λ-) | Casadaban (Casadaban, 1976) |
| JJD14 | *E. coli* MC4100 ($\lambda$ cI857 S<sub>am</sub>7) λIII | Wang (Wang, 2006) |
| JJD459 | *E. coli* MC4100 + pS<sub>wt</sub> | This study |
| JJD251 | *E. coli* MC4100 ($\lambda$ cI857 S<sub>105</sub>/A99V) | Wang |

### Plaque assays

To obtain plaques of lysis-deficient λ phages, an *E. coli* strain with the pSwt (Wang 2006) plasmid, which expresses holin in *trans*, was constructed. This *E. coli* strain was grown overnight in TB broth (5 g NaCl and 10 g tryptone in 1 L water) containing 0.2% maltose and ampicillin (100 μg/ml) at 37°C. The overnight culture was diluted with equal volume of TB + maltose + ampicillin and grown for another 2 h. A 100 μL aliquot of these cells was mixed with phage lysates and incubated at room temperature for 20 min to allow pre-adsorption. This mixture was then added to 3 mL of molten H-top agar (Miller 1992), gently vortexed, and overlaid onto freshly prepared plates containing 35 mL LB agar. The plates were then incubated at 37°C and plaques were counted after 18–22 h. For each time point, single-cell burst sizes from ≈100 cells were estimated.

### Thermal induction and lysis of single cells

Cultures of the lysis-deficient lysogen (*E. coli* MC4100 (λ cI857 S_am_7) λIII) were grown overnight at 30°C in LB broth supplemented with 0.2% glucose (LBG). Overnight cultures were diluted 100-fold in LBG and grown in a 30°C shaking incubator (200 rpm) the cultures reached an OD_600_ of 0.3–0.4. The cultures were transferred to a 42°C shaking water bath for 20 min to induce lysis. Induced cultures were quickly diluted in LBG (prewarmed at 42°C) and 200-μl aliquots were transferred into wells of a 96-well plate such that each well received 0.25 cells/well. This degree of dilution results in mostly empty wells and only 10% of wells will contain more than one cell (Delbrück 1945). The plate was quickly transferred into a prewarmed plate reader (Tecan Infinite M200Pro) and incubated at 37°C with constant agitation (orbital shaking, amplitude 6, frequency 141.9 rpm). After incubation in the plate reader for the required time, a multi-channel pipette was used to quickly transfer 100 μL of chloroform into each well. The plate was then shaken at room temperature for 10 min. This treatment ensured lysis of cells to release phage virions. A 100-μL aliquot of the supernatant aqueous layer was carefully collected and used in plaque assays to enumerate phages. For the lysogen with functional holin, an hour of incubation was sufficient for natural lysis.

## RESULTS AND DISCUSSION

To estimate single-cell burst sizes, we employed a lysis-deficient λ lysogen, which can be thermally induced to initiate the lytic cycle. Once induced, the host cells do not divide, allowing the lytic cycle to proceed unhindered. Thus, cells are “phage factories” that assemble phages until they are chemically lysed using chloroform (Fig. 1). Adsorption of the released phages to host LamB receptors was minimized by supplementing the growth media with glucose, which represses LamB expression. The released phages were lysis deficient, thus an *E. coli* host that expresses functional holin in *trans* was employed for plaque assays. This system enabled us to estimate single-cell burst sizes at different lysis times.

**Fig.1.**
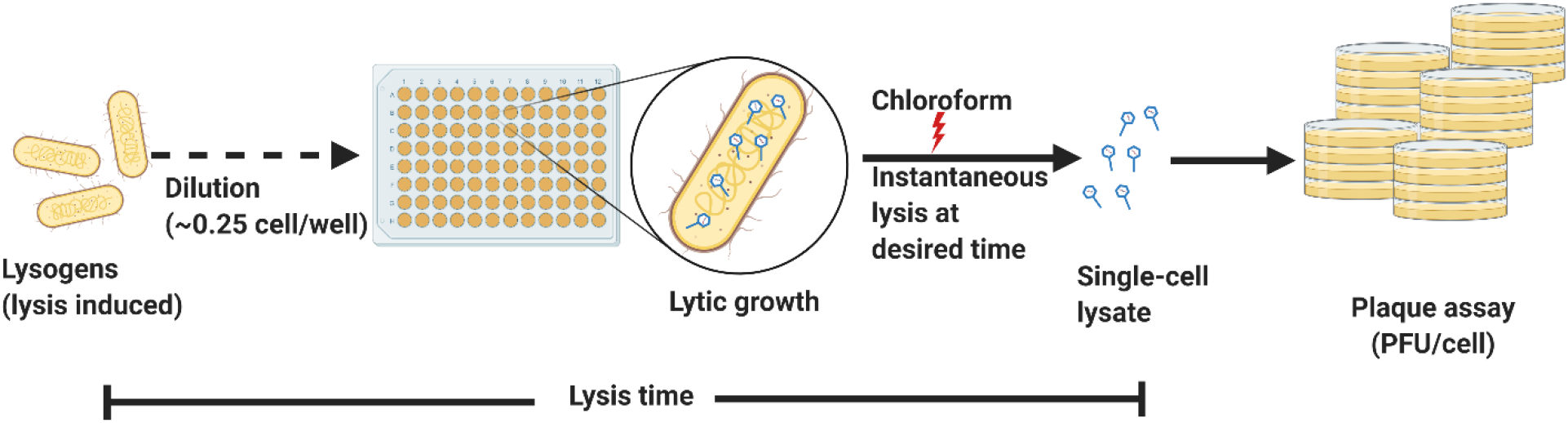
Single-cell lysates can be used to estimate burst size. An induced culture of *E. coli* lysogen with lysis-deficient λ phage was diluted and aliquoted into a 96-well plate to give ≈ 0.25 cells/well, which minimizes the probability of having multiple cells per well. The lytic cycle was allowed to proceed until the cells were chemically lysed, which resulted in single-cell lysates in some wells. The contents of each well were used in plaque assays to estimate burst size.

Induced cells were diluted such that each well of a 96-well plate received 0.25 cells to minimize the probability that wells received multiple cells. If performed accurately, each plate will produce 74 wells with nothing but the media. Plaque assays using the contents of such wells will result in no plaques. As a result, to calculate the burst sizes for at least 100 cells, around 500 plaque assays were performed for each time point. However, batch-to-batch variation resulted in empty wells ranging from 74 to 52 suggesting that the wells received 0.25 to 0.61 cells/well. Based on this range, we can predict that the number of wells that received two cells ranged from 2% to 10% and frequencies for three or more cells are negligible. The burst sizes obtained from wells with multiple cells will lie at the far end of the burst size distribution and might explain the right skewness of these distributions (Fig. 2, upper right panel). The mean burst sizes for approximately 100 cells for each time point show a steady sigmoidal increase with lysis time until 3 h, beyond which no substantial increase is observed. Our data shows that the viral burst size initially accelerates exponentially with the lysis time and then saturates at longer lysis times (Fig. 2, left and bottom right panels). Motivated by this data, we phenomenologically model the burst size (*BS*) as a function of lysis time (*LT*) by the equation

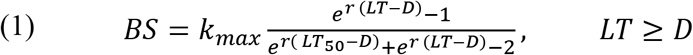

where *D* is a time delay such that *BS* = *0* for *LT* ≤ *D. k_max_* represents the maximum *BS* that can be characterized as the cellular capacity to produce phages. *LT*_50_ is the lysis time where *BS* = *k_max_*/2 and *r* is the exponential growth rate. Fitting eq. (1) to data we estimate *D* ≈ 25 *mins*, *r* ≈ 0.027 *min*^-1^, *LT*_50_ ≈ 119 *mins*, *k_max_* ≈ 1430 *pfu*/*cell* (Fig. 2).

**Fig. 2.**
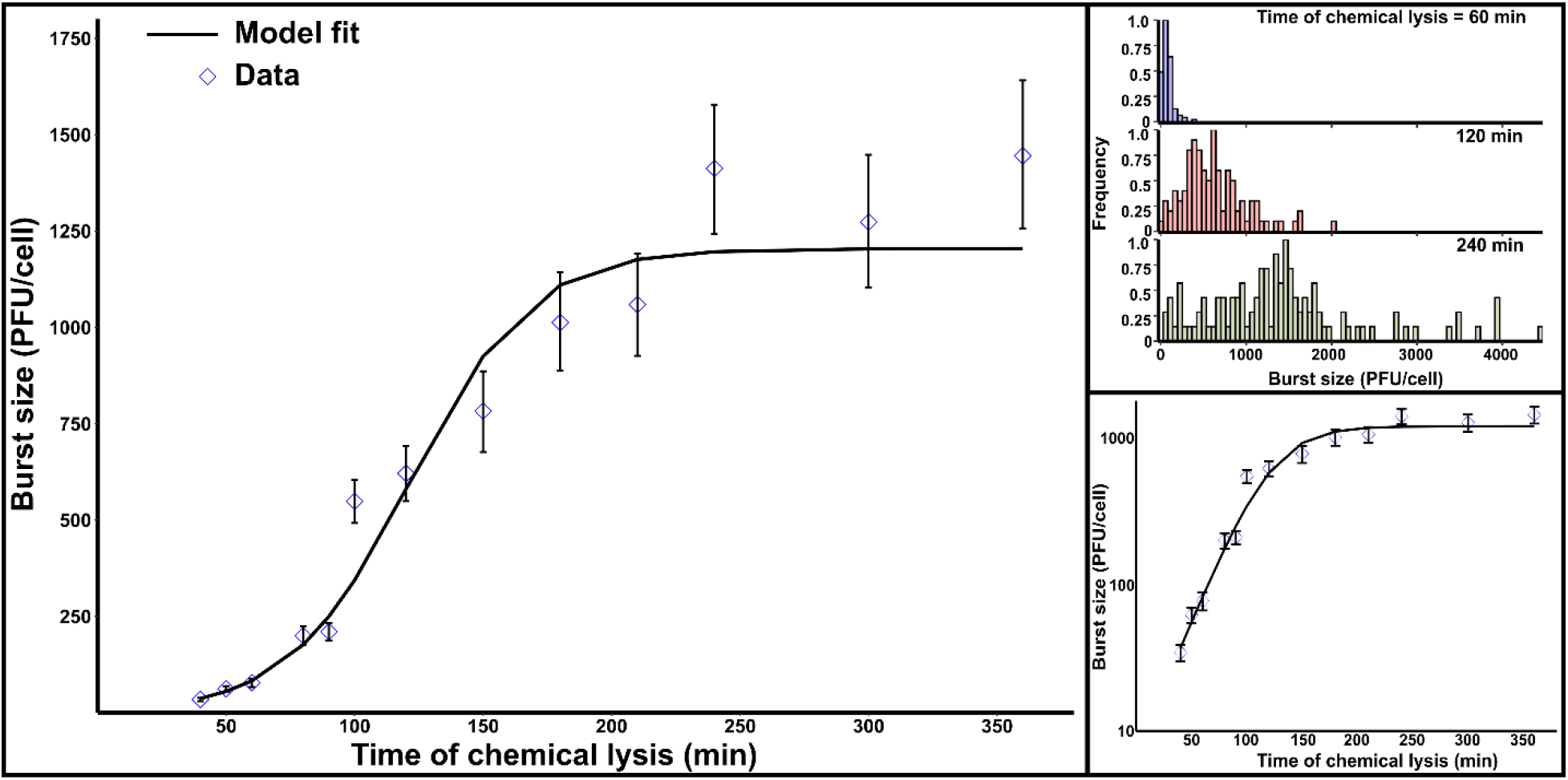
Burst size first accelerates exponentially and then saturates at longer lysis times. After inducing the lytic cycle, single cells of lysogenic *E. coli* with lysis-deficient λ phage were chemically lysed at regular intervals. The mean burst size calculated for approximately 100 cells for each lysis time are shown in the left panel. The error bars represent 95% CIs after bootstrapping (1,000 replicates) and the black line represents the model fit (eqn. (1)). The upper right panel shows the burst sizes distributions for three lysis times. The bottom right panel shows the exponential increase in burst size when the data is plotted on the Y-axis using a log scale.

Next, we investigated if cell-to-cell differences in the lysis time affects variation in the viral burst size. Previous studies using the lysis-deficient lysogen have demonstrated the lytic effect of chloroform on induced cells. When added to an induced culture, chloroform permeabilizes the inner membrane of these lysogens, which results in instant loss of turbidity (Raab *et al*., 1988; Chang, Nam and Young, 1995). Lysis events mediated by functional holin will show cell-to-cell variation in lysis timing. However, chloroform-induced lysis will minimize such variations, especially in our experimental set up where a well most likely contains a single cell (Fig. 1). The use of a multichannel pipette to quickly add chloroform to each well minimizes the variation in lysis timing of single cells. This instantaneous lysis ensures negligible contribution of lysis time variation to the burst size variation.

To quantify the cell-to-cell variation or noise in burst size, we estimated noise as a unitless metric, the coefficient of variation (*CV* = standard deviation divided by the mean). We compared the burst size noise in chemically lysed (using chloroform) cells to that of a naturally lysing holin mutant (see strain JJD51 in Table 1., Kannoly *et al*., 2020) with a mean lysis time of ≈40 min (Fig. 3). If the noise in lysis timing significantly contributes to the noise in burst size, then the burst size *CV* of the naturally lysing holin mutant will be significantly higher when compared to the chemically lysed mutant. We used the R package cvequality Version 0.1.3 (Marwick and Krishnamoorthy, 2019) to test for significant differences in the *CVs*. The noise in burst size estimated from natural and chemical lyses were not significantly different (Asymptotic test, *p* = 0.81; Modified signed-likelihood ratio test, *p* = 0.82) suggesting that the noise in lysis timing does not significantly contribute to the noise in burst size.

**Fig. 3.**
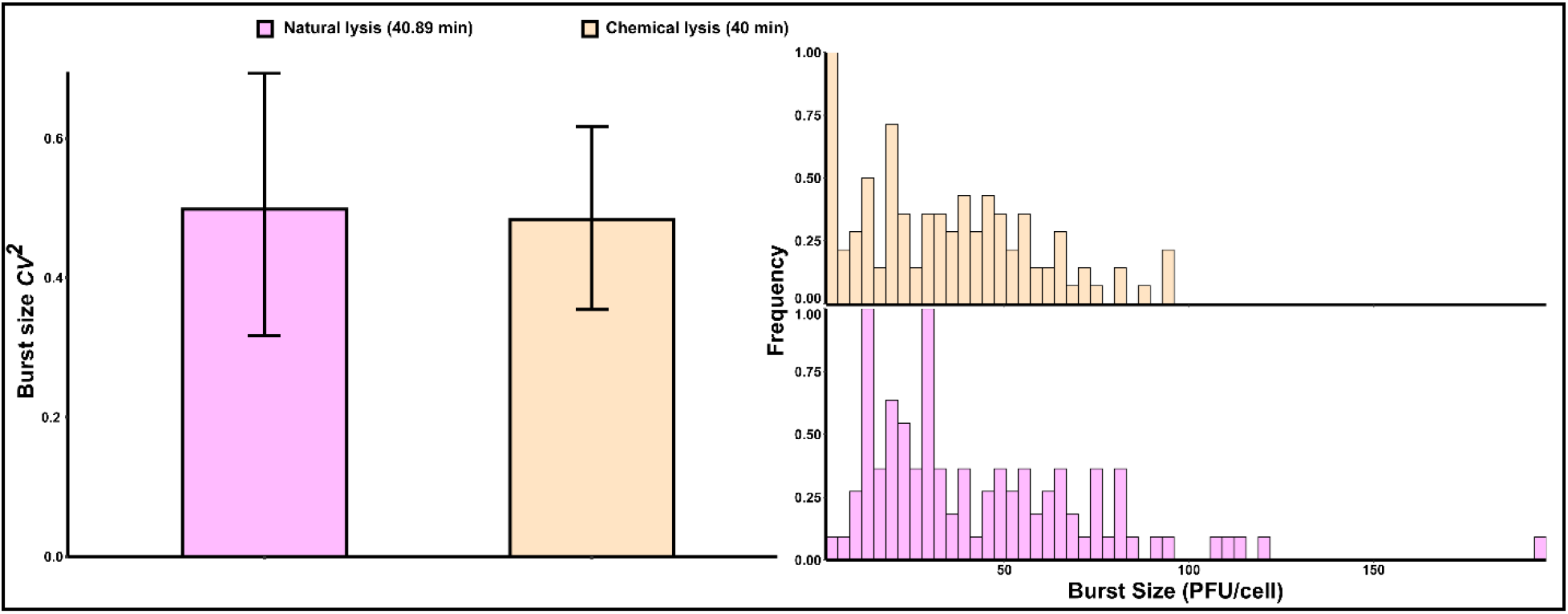
Chemically and naturally lysed cells show a similar cell-to-cell variation (noise) in burst size (*CV^2^*). An induced culture of *E. coli* lysogen with lysis-deficient λ phage was chemically lysed at 40 min to estimate the noise in burst size as quantified using the coefficient of variation squared (*CV^2^*). This noise was compared to that of lysogenic cells harboring a mutant holin that shortens the mean lysis time to 40.89 min (left). The distribution of single-cell burst sizes for both lysogens are shown on the right. Error bars, 95% CIs after bootstrapping with 1,000 replicates.

Using the burst size estimations, we further calculated the noise in burst size at different lysis times. Interestingly, burst size noise appears to remain fairly constant with increasing mean burst size (Fig. 4). Inspired by this data, we attempted to explain the constant burst size noise using the parameters in eqn. (1). The most likely explanation for the experimentally observed constant burst size noise is that cell-to-cell differences in burst size originate from differences in cellular capacity (*k_max_*) to produce phages (see Supplemental information).

**Fig. 4.**
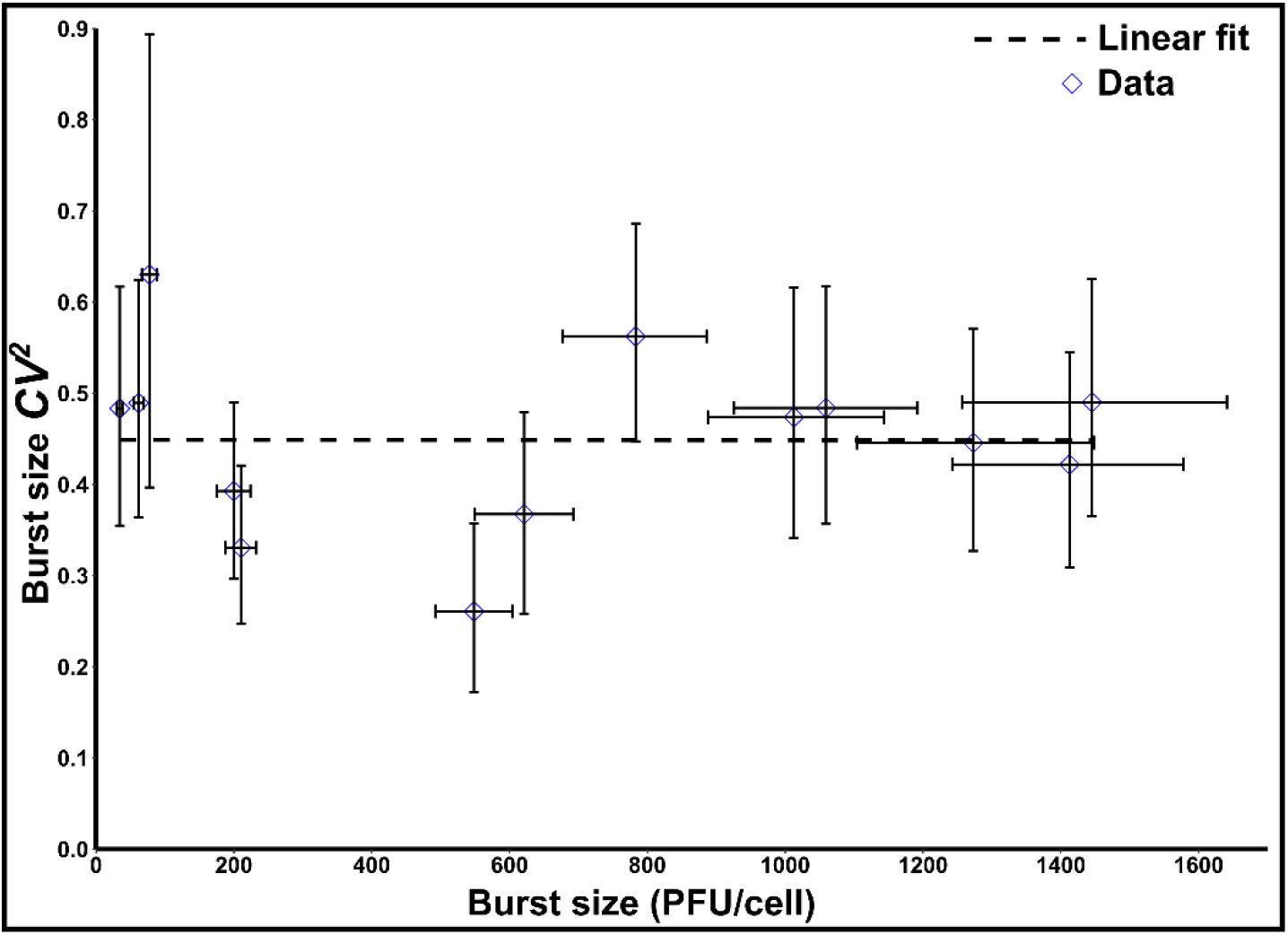
Cell-to-cell variation (noise) in burst size remains constant with increasing mean burst size. Noise (*CV^2^*) in burst size is shown plotted against mean burst sizes estimated for a range of lysis times. The dotted line is a linear fit of the data. Error bars, 95% CIs after bootstrapping (1,000 replicates).

First, we asked if the lack of nutrients is a limiting factor for continuous phage production. A previous study estimated that a T4 phage infection with a burst size of 200 requires 30% of the host energy supply (Mahmoudabadi, Milo and Phillips, 2017). Being smaller than T4 phage, phage λ may be assumed to have similar or lower energy requirements. *E. coli* growing in a complex and rich medium such as LB has a carrying capacity of ≈ 5 × 10^9^ CFU/mL so a single metabolically active cell in 200 μL of LB can theoretically saturate the medium with a total of about a billion cells. Yet the burst size in a single cell only reaches a maximum of thousand phages indicating that assembly and maturation of virions is not limited by nutrients required to sustain cell metabolism.

Another possibility is that phage production is limited by the available intracellular space. The estimated volume of an *E. coli* cell is ≈ 4.4 μm^3^ (Volkmer and Heinemann, 2011) and that of T4 phage is ≈ 3.54 × 10^-4^ μm^3^ (Nifong and Gillooly, 2016), which suggests that an *E. coli* cell can theoretically harbor ≈ 10^4^ T4 virions. However, it is estimated that only 18% of the cell volume can be occupied by proteins (17%) and DNA (1%). The rest of the cell’s volume is occupied by water (70%), ribosomes (8%), and other miscellaneous macromolecules (Guo *et al*., 2013; Sajed *et al*., 2016). If we assume that the 18% of cellular volume occupied by proteins and DNA (0.792 μm^3^) can be entirely replaced by T4 virions, then the maximum T4 burst size can reach ~2,200 virions. Moreover, despite phages redirecting cell resources from replicating cell DNA and proteins to replicating phage DNA and proteins, this redirection cannot be 100% efficient. That is, cells still need to maintain housekeeping functions to survive, thus the available space for phages must be somewhat less than 18% of the cell’s volume.

To further complicate matters, we observed that cells continue to grow after induction of the lytic cycle, although this growth slows towards the end of the lytic cycle (Fig. S1). As burst size is positively correlated to the cell volume (Fig. S2), we suspect that the cellular capacity (*k_max_*) for phage production is mostly constrained by the available intracellular space, especially as the burst size approaches saturation at ~ 1000 virions at 3 h. Thus, the differences in cellular capacity to produce phages might be explained by the observed differences in cell size.

To test this idea, we used fluorescence assisted cell sorting (FACS) to estimate the burst size of cells that fell within a narrow size window. If cell size is the sole determinant of cellular capacity, then burst size noise estimated from cells of similar size would approach zero. However, the burst size noise (*CV*) estimated from similar-sized cells with a mean burst size of 82.62 was 0.42 (Fig. S3). These numbers are comparable to burst size measurements of the same strain by standard techniques reported by another study, where mean burst size was 88.5 with a *CV* of 0.80 (Wang 2006). However, this study estimated burst size averaged across a phage population, which might explain the higher noise when compared to single cell estimations. Moreover, the lower *CV* for the FACS-sorted cells probably reflects the narrower range of cell sizes for sorted cells compared to those found in ordinary *E. coli* broth cultures. This finding suggests that, in addition to cell size, there are other factors that determine the cellular capacity to produce phages.

What other factors might influence phage production? Being obligate intracellular parasites, phages use the cellular machinery for genome replication and assembly of virions. The concentrations of ribosomes and other proteins, which play key roles in the translation and replication processes, depend on the physiological and metabolic states of a given cell. For example, the distribution of ribosomes in newly divided daughter cells are unequal, and the number of ribosomes start to increase half way through the cell cycle and peaks close to cell division (Chai *et al*., 2014). For our burst size estimations, we induced exponentially growing cells that are not synchronized with respect to their cell cycles. Cells at different stages of growth may present with distinct environments affecting the phage life cycle. Therefore, it is not hard to imagine that the intracellular milieu of a cell would have profound effects on overall phage production. A study using T4 phages showed that phage productivity in cells close to cell division was almost three times greater that the productivity of newly divided daughter cells. The study also found that the intracellular RNA levels and not the DNA levels were strongly correlated to phage productivity (Storms *et al*., 2014). It is interesting to note that following induction of the lytic cycle, cytokinesis is blocked by λ via ZipA-dependent inhibition of FtsZ (Haeusser et al., 2014). It is possible that as the lytic cycle proceeds, the increased availability of intracellular resources accelerates phage production. It would be interesting to explore how cell cycle synchronization, which can be effected with serine hydroxymate, impacts burst size and lysis time noise. In addition, experiments where relative concentrations of key cellular proteins are simultaneously quantified along with phage burst size may provide insights into the factors affecting phage production capacity.

In this study, we showed that it was possible to simultaneously estimate phage burst size and variation in burst size at different lysis times. By titering the phages prior to and after one round of infection and lysis, a one-step growth curve can only estimate lysis time and burst size values averaged across a phage population. The one-step growth curve method cannot be used to estimate the cell-to-cell variation in burst sizes. Burnet invented the first method for single-cell burst size estimation by diluting a phage-infected suspension of bacterial cells in to small aliquots such that each aliquot contained on an average less than one infected bacterium (Burnet, 1929). This strategy ensured that only a small fraction of aliquots would contain more than one infected bacterium. After incubation for sufficient time to allow lysis of all bacteria, plaque assays revealed the distribution of burst sizes.

This method was later improved to estimate the burst size distribution in a larger sample size (Delbrück, 1945). These early methods used free phages to initiate infection by allowing adsorption onto growing cells and required quick dilution and distribution of cells before the first burst occurred. By contrast, our method uses a lysis-deficient lysogen, which allowed us to induce the lytic cycle simultaneously in all cells as well as to lyse the cells at desired times. This allowed more accurate estimations of burst size mean and its distribution at different lysis times. Using this method, we observed that the noise in lysis timing does not significantly contribute to the noise in burst size and that the burst size noise remained constant across different lysis times. We surmise that the likely explanation for the experimentally observed constant burst size noise is that cell-to-cell differences in burst size derive from differences in cellular capacity (*k_max_*) to produce phages (see Supplemental information).

Cell-to-cell variation in lysis timing estimated for a collection of mutants with a wide range of mean lysis times was considerably lower than the variation in burst size at different lysis times (Kannoly *et. al*., 2020). A high degree of variation in burst size at different lysis times may have consequences for phage λ’s fitness. Phage fitness is highly correlated with the host physiological state, which in turn is dependent on the environment. In studies that used one-step growth curves, it was observed that burst size increases and/or lysis time shortens as the physiological state of the host improves (Webb, Leduc and Spiegelman, 1982; Abedon, 1989; Kokjohn, Sayler and Miller, 1991; Proctor, Okubo and Fuhrman, 1993; Middelboe, 2000; Abedon, Hyman and Thomas, 2003; Gnezda-Meijer *et al*., 2006; Birch, Ruggero and Covert, 2012; Golec *et al*., 2014). Such short-term changes in the values of phage traits triggered by changes in the host is described as viral phenotypic plasticity (Hadas *et al*., 1997; Abedon, Herschler and Stopar, 2001; You, Suthers and Yin, 2002; Zheng *et al*., 2008). A recent theoretical approach has suggested that burst size plasticity drives ecological and evolutionary dynamics by strengthening dynamic feedbacks between a phage, its host, and the environment (Choua and Bonachela, 2019). It was hypothesized that plasticity in burst size is more important than the plasticity in lysis time especially under favorable growth conditions, which allows production of more virions within a shorter lytic cycle. Plasticity provides phenotypic diversity in a phage population and may render sensitivity to the host environments without the need for genetic changes. Our findings revealed a high degree of variation in burst size, which conform to these theoretical predictions.

## ACKNOWLEDGEMENTS

This work was made possible by grant number 1R01GM124446-01 from the National Institutes of Health. We thank Ing-Nang Wang for stimulating conversations about phage λ and for the phage and bacterial strains described herein. We also thank the members of the Dennehy Lab for support, advice, discussions, and feedback.

